# Ion exchange biomaterials to capture daptomycin and prevent resistance evolution in off-target bacterial populations

**DOI:** 10.1101/2022.06.10.495716

**Authors:** Shang-Lin Yeh, Naveen Narasimhalu, Landon G. vom Steeg, Joy Muthami, Sean LeConey, Zeming He, Mica Pitcher, Harrison Cassady, Valerie J. Morley, Sung Hyun Cho, Carol Bator, Roya Koshani, Robert J. Woods, Michael Hickner, Andrew F. Read, Amir Sheikhi

**Affiliations:** Department of Chemical Engineering, The Pennsylvania State University, University Park, PA 16802, USA; Department of Biology and Entomology, The Pennsylvania State University, University Park, PA 16802, USA; Department of Chemistry, The Pennsylvania State University, University Park, PA 16802, USA; Department of Materials Science and Engineering, The Pennsylvania State University, University Park, PA 16802, USA; Huck Institutes of the Life Sciences, The Pennsylvania State University, University Park, PA 16802, USA; Department of Internal Medicine, University of Michigan, Ann Arbor, MI 48109, USA; Department of Biomedical Engineering, The Pennsylvania State University, University Park, PA 16802, USA; NTx, 7701 Innovation Way, NE Rio Rancho, NM 87144

**Keywords:** antibiotic stewardship, daptomycin, ion exchange, cholestyramine, biomaterial

## Abstract

Daptomycin (DAP), a cyclic anionic lipopeptide antibiotic, is among the last resorts to treat multidrug resistant (vancomycin resistant *Enterococcus faecium* or methicillin resistant *Staphylococcus aureus*) Gram-positive bacterial infections. DAP is administered intravenously and biliary excretion results in the introduction of DAP (∼5-10 % of the intravenous DAP dose) arriving in the gastrointestinal (GI) tract where it drives resistance evolution in off-target populations of *Enterococcus faecium* bacteria. Previously, we have shown that the oral administration of cholestyramine, an ion exchange biomaterial (IXB) sorbent, prevents DAP treatment from enriching DAP-resistance in populations of *E. faecium* shed from mice. Here, we engineer the biomaterial-DAP interfacial interactions to uncover the antibiotic removal mechanisms. The IXB-mediated DAP capture from aqueous media was measured in both controlled pH/electrolyte solutions and in simulated intestinal fluid (SIF) to uncover the molecular and colloidal mechanisms of DAP removal from the GI tract. Our findings show that the IXB electrostatically adsorbs the anionic antibiotic via a time-dependent diffusion-controlled process. Unsteady-state diffusion-adsorption mass balance describes the dynamics of adsorption well, and the maximum removal capacity is beyond the electric charge stoichiometric ratio because of DAP self-assembly. This study may open new opportunities for optimizing cholestyramine adjuvant therapy to prevent DAP resistance, as well as designing novel biomaterials to remove off-target antibiotics from the GI tract.

**TOC:** 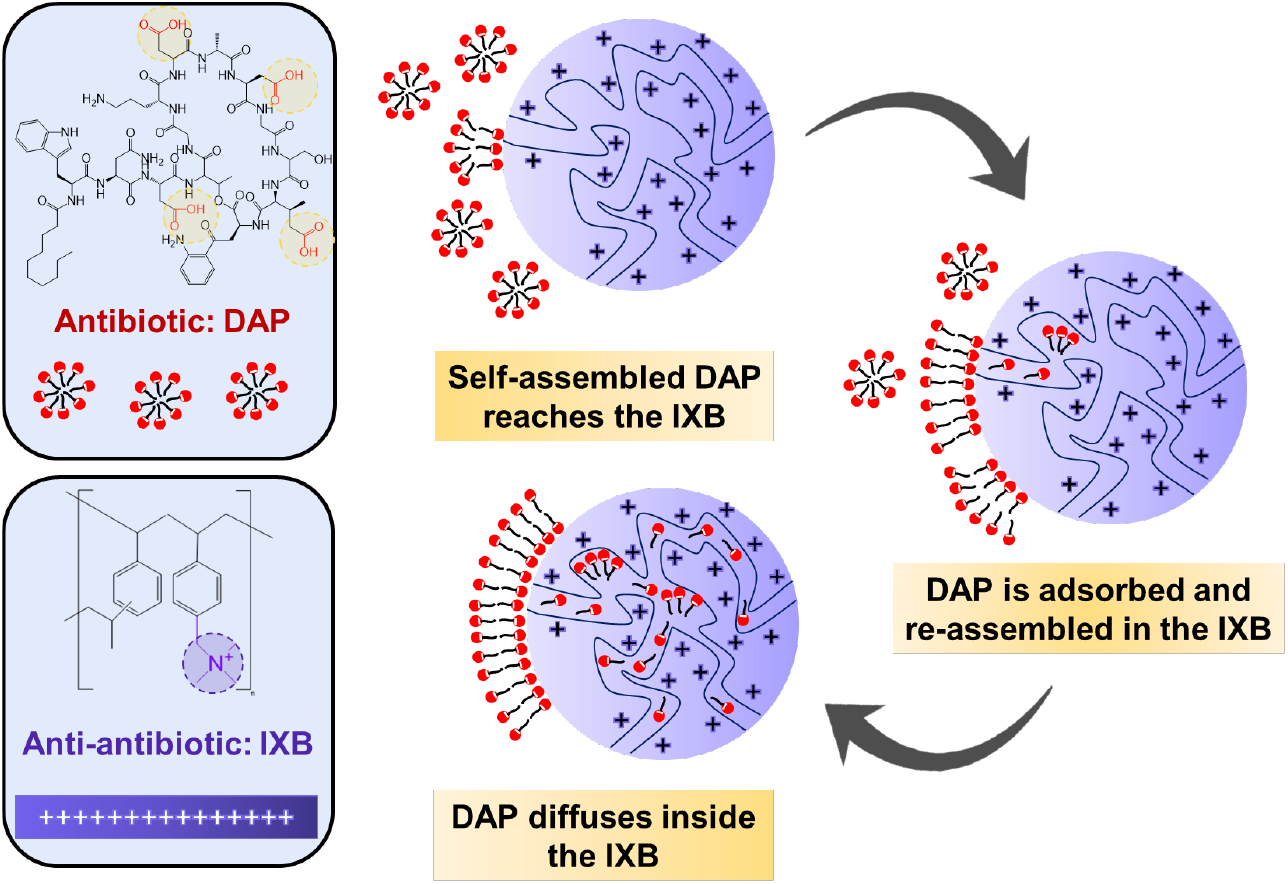

## 1. Introduction

Daptomycin (DAP) is an important first-line antibiotic for the treatment of multidrug resistant Gram-positive bacteria like vancomycin resistant *Enterococcus faecium* (VRE*fm*), a leading cause of hospital acquired infections.^1^ The excellent therapeutic capability of DAP is currently threated by increasing frequencies of DAP resistance in VRE*fm* worldwide.^2, 3^ *E. faecium* normally asymptomatically colonizes gastrointestinal (GI) tracts; transmission is fecal-oral.^4^ GI colonization is a key risk factor for clinical infections,^5^ such as blood stream infections or endocarditis, which can be life threatening.^6^

DAP is administered intravenously (IV) to treat VRE*fm* and other Gram-positive blood stream infections. Most DAP is excreted via urine, but 5-10 % enters the GI tract through biliary excretion, where it has no therapeutic value.^1^ We have found that DAP resistance was present in GI VRE*fm* populations in patients who had received IV DAP therapy, and not in case-matched patients who received a different antibiotic.^7^ This is consistent with off-target selection for resistance, where antibiotic use causes resistance evolution in a population of bacteria that are not the therapeutic target. Direct experimental evidence of off-target selection comes from mouse models: systemic DAP treatment enriched for DAP-resistant VRE*fm* in the GI tract and up-selected a variety of *de novo* resistance mutations that confer DAP resistance.^8, 9^ This raises the prospect that inactivation of DAP in the GI tract would prevent the evolution of DAP resistance without interfering with the capacity of DAP to treat bloodstream infections. This would reduce the risk that patients would acquire DAP-resistant VRE*fm* from their carriage populations in their own GI tract, as well as prevent the onward transmission of DAP resistance in this important hospital-acquired infection. Capture of GI DAP would also prevent DAP-induced perturbations of GI microbiota that are associated with the expansion of populations of other disease-causing infections such a *Clostridium difficile*, and many other disease states associated with the microbiome.^10^

DAP is a cyclic lipopeptide antibiotic, which is produced via fermentation involving decanoic acid-spiked *Streptomyces roseosporus* growth media.^11^ DAP is amphiphilic, composed of 13 amino acids forming a hydrophilic head connected to a decanoyl fatty acid as a lipophilic tail.^12^ It also contains four carboxyl groups, which bear pH-dependent charge. Although the mechanism of the DAP-mediated Gram-positive bacterial cell death is poorly understood,^13^ it is believed that the binding between DAP and the target bacterial cell membrane is associated with bacterial death.^13^ Mediated by hydrophobic interactions between the lipid chain of DAP and phospholipid cell membrane, DAP is inserted into the bacterial cell membrane, depolarizing it, and compromising the intracellular components.^14^ In addition, the antibacterial action of DAP depends on calcium ion-mediated aggregation of hydrophilic head, leading to conformational changes that induce DAP interactions with the bacterial membrane.^15^

Inactivating DAP in the intestines without reducing DAP plasma concentrations would enable the IV use of DAP to eliminate bacteria in the infection sites without driving resistance in the GI tract populations. We have previously shown that feeding mice with an ion exchange biomaterial (IXB) sorbent, cholestyramine, reduced DAP-induced enrichment and shedding of DAP-resistant VRE*fm* by 80-fold^16^ and completely prevented the emergence and shedding of *de novo* resistance mutations. ^8, 9^ However, the mechanism by which this oral adjuvant prevents DAP activity is unclear.

In this work, we aim to uncover and engineer the IXB-mediated DAP capture mechanisms via conducting *in vitro* antibiotic removal experiments in aqueous media with controlled pH, ion types, ionic strengths, bile salt, and phospholipid content as well as in simulated intestinal fluid (SIF). Additionally, the time-dependent antibiotic removal will be modeled to identify the role of adsorption and diffusion. The effect of IXB size on the removal efficacy is further investigated to unravel the effect of diffusion in the sequestration process. Understanding these foundations may help rationally design the next generation of antibiotic sorbents to prevent antimicrobial resistance, as well as help optimize the use of cholestyramine itself as an oral adjuvant therapy for resistance prevention.

## 2. Materials and methods

### 2.1. Materials

Daptomycin (DAP, > 94 %) was purchased from Tokyo Chemical Industry, Japan. Sodium chloride (NaCl, > 99.5 %), calcium chloride dihydrate (CaCl_2_⋅2H_2_O, for molecular biology, ≥ 99.0 %), sodium hydroxide (NaOH, ACS Reagent, > 97 %), hydrochloric acid (HCl, ACS reagent, 37 %), L-α-Lecithin (a concentrate of soybean lecithin consisting of more than 94 wt% phosphatidylcholine and less than 2 wt% triglyceride), phosphate buffer (NH_4_H_2_PO_4_), acetonitrile (CH_3_CN, HPLC grade, > 99.9 %),cellulose acetate centrifuge tube filters (pore size = 0.22 μm), cholestyramine resin (IXB, Dowex® 1×2 Cl^-^ Form), and polymer resin AmberChrom 1×4 (AC4) were purchased from Sigma-Aldrich, USA. Maleic acid (C_4_H_4_O_4_, > 98 %) was purchased from Beantown Chemical Corporation, USA. Sodium taurocholate (C_26_H_44_NNaO_7_S) was procured from Spectrum Chemical, USA. Fasted state simulated intestinal fluid (FaSSIF-V2) powder was supplied from Biorelevant, UK. Milli-Q water with a resistivity of 18.2 mΩ cm was generated from the deionized water passing through an ultrafilter (Biopak Polisher, Millipore, USA). Cation-adjusted Mueller Hinton II Broth (BD Difco) was used as a bacterial culture medium and was purchased from Becton, USA. *Enterococcus faecium* (*E. faecium*), strain BL00239-1, was obtained from a blood stream infection in a patient being treated at the University of Michigan hospital.^9^

### 2.2. Methods

#### 2.2.1 Cryogenic transmission electron microscopy (cryo-TEM)

DAP dispersion, IXB suspension, and DAP-IXB suspension were vitrified using a Vitrobot (Thermo Fisher Scientific, USA), for which the chamber was preconditioned to 4°C and 100 % relative humidity. Holey carbon Quantifoil grids (2 µm diameter with an interspace of 2 µm, Quantifoil Micro Tools GmbH, Germany) were prepared by glow discharge (easyGlow® System, Pelco, USA), to which 3.5 μL of samples (0.1 w/v %) were deposited and blotted, immediately followed by plunging into liquid ethane. Cryo-TEM images were acquired using a Talos Arctica TEM (Thermo Fisher Scientific, USA), which was equipped with Falcon 4 Direct Electron Detector (Thermo Fisher Scientific, USA) and controlled by EPU software (Thermo Fisher Scientific, USA, version 2.12.0.2771REL). Imaging conditions were as follows: 200 kV; 57,000x in magnification; counted mode; total dose = 15 e/A^2^; nano probe; spot size = 3; and C2 aperture = 50.

#### 2.2.2 Preparation of IXB with different particle sizes

Polymer resin AC4s were milled to obtain different particle size distribution samples using a CryoMill (Retsch, Germany). Dry resins did not mill well, so samples were exposed to ambient air for a minimum of 2 h to absorb moisture before grinding. The samples were then milled using a steel ball, at milling frequencies varied between 20 Hz to 30 Hz, and milling times ranging from 30 s to 8 min. The parameters were adjusted to provide different particle size distributions, which were then characterized by scanning electron microscopy (SEM, ThermoFisher Verios G4, USA). After milling, the resins were soaked in a NaCl solution (3 M) overnight. The resins were then washed three times with deionized water, including an overnight soak, and then air-dried under fume hood. Particle size distribution of resin was determined by SEM imaging. After imaging, particle size was determined by the ImageJ software (version Java 1.53e).^16^

#### 2.2.3 Simulated intestinal fluid (SIF) preparation

To mimic intestinal fluid *in vitro*, fasted state simulated intestinal fluid (FaSSIF) and fed state simulated intestinal fluid (FeSSIF) were prepared based on standard protocols.^17^ Sodium taurocholate was selected as cholic acid is among the most common bile acids in human bile.^18^ To prepare the FaSSIF, 1.392 g of NaOH pellets, 2.22 g of maleic acid, and 4.01 g of NaCl were dissolved in 0.99 L of Milli-Q water. Then, the pH of solution was adjusted to 6.5 using a NaOH solution (0.1 M), and the total volume was increased to 1 L with Milli-Q water. The solution was then added to 1.79 g of FaSSIF-V2 powder and stirred for 1 h at room temperature. To prepare the FeSSIF solution, 8.25 g of sodium taurocholate was added to 250 mL of FaSSIF solution and stirred at room temperature to completely dissolve the sodium taurocholate. Then, 2.95 g of lecithin was added and continued stirring for 4 h to from a clear solution. The final volume was adjusted to 1 L with the rest of FaSSIF solution.

#### 2.2.4 DAP removal

The DAP removal experiments were conducted in a batch process. DAP stock solutions were prepared by dissolving 100 mg of DAP in 5 mL of Milli-Q water (concentration = 20 mg mL^-1^). Then, DAP solutions with varying concentrations ranging from 1 mg mL^-1^ to 20 mg mL^-1^ were prepared by the successive dilution of stock solution with Milli-Q water. The solution pH was adjusted to 6.5 by adding a NaOH solution (0.5 M). The IXB (cholestyramine or AC4, 8 mg) were added to the DAP solution, followed by vortexing for 5 min and placing the vial on the nutating mixer (Fisherbrand, USA) to agitate at 60 rpm for a desired incubation time depending on the type of experiments. The samples were centrifuged at 5000 ×**g** for 5 min, and the supernatant was collected and assessed using a UV-vis spectrophotometer (Tecan Model Infinite 200 Pro, USA) at λ_max_= 364 nm to measure the unabsorbed DAP concentration.

#### 2.2.5 UV-vis spectroscopy for DAP concentration measurement

To measure the DAP concentration, calibration curves were obtained for each experimental condition (14 calibration curves, shown in **Figure S1**) by recording the absorbance of DAP solutions with predetermined concentrations ranging from 0.05 mg mL^-1^ to 20 mg mL^-1^ at λ_max_= 364 nm using the UV-vis spectrophotometer. The kynurenine residue in DAP causes the absorbance peak at 364 nm ^19^. The calibration curves were used to determine the concentration of DAP in the supernatant after adsorption by the IXB.

#### 2.2.6 High-performance liquid chromatography (HPLC) for DAP concentration measurement

The HPLC (HP 1000, Thermo Fisher, USA) equipped with UV detector set at 224 nm was used to measure DAP concentrations below 50 mg L^-1^. The mobile phase consisted of ammonium phosphate buffer (NH_4_H_2_PO_4_) (40 mM, pH 4.0), and acetonitrile/water (10:90 v/v). Chromatographic separation was achieved using an Agilent ZorbaxTM C_18_ analytical column (Length: 150 mm, inner diameter: 4.6 mm, particle size: 5 µm) (AgilentTM, USA). The volume of the injection was 20 µL and the flow rate was 0.5 mL min^-1^. Column temperature was maintained at 25 °C. The peak areas detected at 224 nm were defined as analytical signs, with detection concentration varying from 5 mg L^-1^ to 200 mg L^-1^ set as calibration line **(Figure S2**).

#### 2.2.7 Antibiotic removal kinetics

To investigate the DAP adsorption kinetics, DAP solutions with a concentration of 8 mg mL^-1^ were prepared (pH = 6.5 adjusted using 0.5 M NaOH, total volume 1 mL). Then, IXB (8 mg) was added to them, followed by vortexing for 5 min and placing them on the nutating mixer to agitate at 60 rpm for up to 24 h. At each time point, the samples were centrifuged at 5000 ×**g** for 5 min, and the supernatant was collected and assessed using the UV-vis spectrophotometer at λ_max_= 364 nm to measure the DAP concentration in the supernatant. The DAP removal capacity of IXB was calculated based on equation (Eqn.) 1:

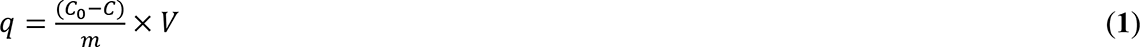

where *q* (mg g^-1^) denotes the mass of DAP adsorbed per 1 g of IXB at time *t*, *m* is the mass of IXB (g), *C*_0_ (mg mL^-1^) is the initial concentration of DAP, and *C* (mg mL^-1^) is the concentration of DAP in the supernatant at time *t*.

#### 2.2.8 Equilibrium adsorption

Equilibrium batch adsorption measurements were performed to obtain the maximum DAP removal capacity of IXB. One mL of DAP solution with desired concentrations ranging from 1 to 20 mg mL^-1^ were prepared (pH = 6.5 adjusted by 0.5 M NaOH), and 8 mg of IXB was added to them. The solutions were then vortexed for 5 min and placed on the nutating mixer with a 60 rpm agitation speed for 4 h. The samples were centrifuged at 5000 ×**g** for 5 min, the supernatants were collected and assessed using the UV-vis spectrophotometer at λ_max_= 364 nm to measure DAP concentration. Removal percentage (*R,* %) and equilibrium removal capacity (*q_e_,* mg mL^-1^) were calculated using Eqns. 2 and 3, respectively:

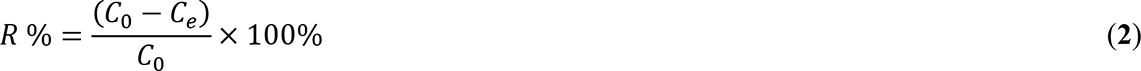

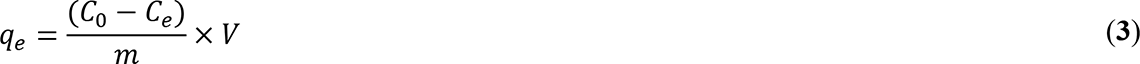

where *C_0_* (mg mL^-1^) denotes initial DAP concentration, *C_e_* (mg mL^-1^) is the equilibrium DAP concentration after removal, *m* (g) is the IXB mass, and *V* (mL) represents the total volume of solution.

#### 2.2.9 Effect of pH on the antibiotic removal

The IXB-mediated removal of DAP (8 mg mL^-1^) at pH ∼ 1.2 - 12 was investigated to examine the effects of functional group ionization on IXB-DAP interactions. The pH of DAP solutions was adjusted by adding HCl (0.5 M) or NaOH (0.5 M) solutions to the DAP stock solution. Then, 8 mg of IXB was added to the DAP solutions with varying pH, followed by vortexing for 5 min and placing them on the nutating mixer with 60 rpm agitation speed for 4 h. The DAP concentration was measured by collecting the supernatant after centrifugation at 5000 ×**g** for 5 min, followed by measuring using the UV-vis spectrophotometer at λ_max_= 364 nm. The removal percentage and capacity were calculated based on Eqns. 2 and 3, respectively.

#### 2.2.10 Effect of ionic strength and ion type on the antibiotic removal

To investigate the effect of monovalent and divalent ions on the IXB-DAP interactions, the IXB-mediated removal of DAP (8 mg mL^-1^) at varying NaCl and CaCl_2_ concentrations ranging from 10 mM to 500 mM was investigated. In this experiment, the pH of all DAP solutions was maintained constant at 6.5 using a NaOH solution (0.5 M). Eight mg of IXB was added to the DAP solutions containing varying concentrations of sodium (Na^+^) or calcium (Ca^2+^) ions. The solutions were then vortexed for 5 min and placed on the nutating mixer with 60 rpm agitation speed for 4h. The DAP concentration in the supernatant was quantified by collecting the supernatant after centrifugation at 5000 ×**g** for 5 min and measuring the absorbance using the UV-vis spectrophotometer at λ_max_=364 nm. The removal percentage and capacity were calculated using Eqns. 2 and 3, respectively.

#### 2.2.11 Effects of SIF components on the antibiotic removal

The IXB-mediated removal of DAP (8 mg mL^-1^) at varying concentrations of maleic acid (5-100 mM), bile acid (1-12 mM), and lecithin (0.5-4 mM) was determined to investigate the influence of SIF components on the IXB-DAP interactions. For these experiments, the pH of DAP solutions was adjusted to 6.5 using a NaOH solution (0.5 M). Eight mg of IXB was added to the DAP solutions containing each of the SIF components at varying concentrations. The solutions were then vortexed for 5 min and placed on the nutating mixer with 60 rpm agitation speed for 4h. The DAP concentration was measured by collecting the supernatant after centrifugation at 5000 ×**g** for 5 min, followed by recording the absorbance using the UV-vis spectrophotometer at λ_max_=364 nm. The removal percentage and capacity were calculated using Eqns. 2 and 3, respectively.

#### 2.2.12 Fourier-transform infrared (FTIR) spectroscopy

The functional groups of DAP, IXB, and DAP-IXB complex were identified using a FTIR spectrometer (ThermoFisher, Pleasantville, NY), equipped with a Diamax attenuated total reflectance (ATR) accessory. The samples were dried in an oven overnight at 50°C to remove water. Samples were placed directly onto the ATR crystal fixed at an incident angle of 45°, and maximum pressure was applied by lowering the tip of pressure clamp. The recorded spectra were averaged from a total of 500 scans at transmission mode ranging from 4000 cm^-1^ to 500 cm^-1^ and a resolution of 6 cm^-1^.

#### 2.2.13 Hydrodynamic size measurement by dynamic light scattering (DLS)

The hydrodynamic size of DAP was measured using DLS (Malvern Zetasizer Nano series, UK) at 90^°^ scattering angle and ambient temperature based on the *z*-average (cumulants mean) of intensity measurements. The concentration of DAP stock solution was adjusted to 0.1 w/v % using Milli-Q water. Then, the solutions were transferred to low-volume quartz cuvettes (ZEN2112) for conducting the measurements.

#### 2.2.14 ζ-potential measurement by electrophoretic light scattering (ELS)

The ζ-potential of DAP was determined by measuring the electrophoretic mobility using Nano ZS Zetasizer (Malvern Instrunments, UK). For measuring the ζ-potential, the DAP concentration was first adjusted to 0.1 w/v % by diluting the DAP stock solution (2 w/v %) using Milli-Q water. The solution pH was adjusted to 6.5 by adding NaOH (2 M), and then the solution was transferred to the universal dip cell kit. Since the DAP size was > 200 nm and the final electrolyte concentration was > 1 mM, the κ value was in the order of 1-10 nm, and the κ*a* was >> 100 (κ is the Debye-Hückel parameter, and *a* is the radius of the particle), thus the ζ-potential can be calculated by applying electrophoretic mobility to Smoluchowski equation.^20^

#### 2.2.15 Evaluating antibiotic activity of uncaptured DAP using broth microdilution assays

Broth microdilution assays were used to directly quantify the effects of IXB on the antibiotic activity of DAP against patient-driven *Enterococcus faecium* (*E. faecium*). Antibiotic capture was performed in Milli-Q water at a pH of 6.5 prior to centrifugation at 16,300 ×**g** for 5 min and passage of the supernatant through a cellulose acetate filter (pore size = 0.22 μm). All assays were conducted in accordance with the guidelines of the Clinical Laboratory Standards Institute,^21^ while using a previously isolated DAP-susceptible BL00239-1-S (Minimum Inhibitory Concentration (MIC) = 2.1 μg mL^-1^) isolate.^9^ Initial DAP concentrations (*i.e.,* the DAP concentration prior to antibiotic capture) were used to generate the 2-fold serial DAP dilutions. After incubation at 35°C for 24 h, bacterial cell densities were measured using a plate reader (BioTek Synergy H1 Plate reader, Agilent Technology, USA) at an optical density of 600 nm. The resulting optical density values were fitted to a Hill function as described previously,^7^ with reductions in initial DAP concentration resulting in a right-shift in the growth curve.

## 3. Results and discussion

### 3.1 IXB-mediated DAP removal kinetics

Figure 1a presents the chemical structure of DAP and IXB respectively. To understand the interactions between DAP and IXB, the morphology of DAP, IXB, and the DAP-IXB complex was analyzed using cryo-TEM. As shown in Figure 1b, DAP formed sphere-like self-assembled particles with an average diameter of 112 ± 31 nm (68 particles), which is a result of hydrophobic tail-mediated micelle formation.^22^ The formation of such spherical micelles or aggregates have been reported at DAP concentrations higher than the critical micelle concentration CMC ∼ 0.147 mg mL^-1^.^22^ Since the average size of IXB (cholestyramine) is around 20 μm (as examined using an optical microscope shown in **Figure S3**), only the edge of IXB was imaged by cryo-TEM. The purple dashed lines in Figure 1b show the boundary of IXB. At the edge of DAP-IXB complex, layers of electron-dense materials (shown with red arrows) were observed, which may be attributed to the DAP layer at the interface. When DAP micelles contact a solid substrate, they may fuse together and eventually rupture to form a supported lipid bilayer (SLBs) on IXB. It has been shown that the spreading of amphiphilic molecules on solid supports may yield SLBs.^23, 24^

**Figure 1.**
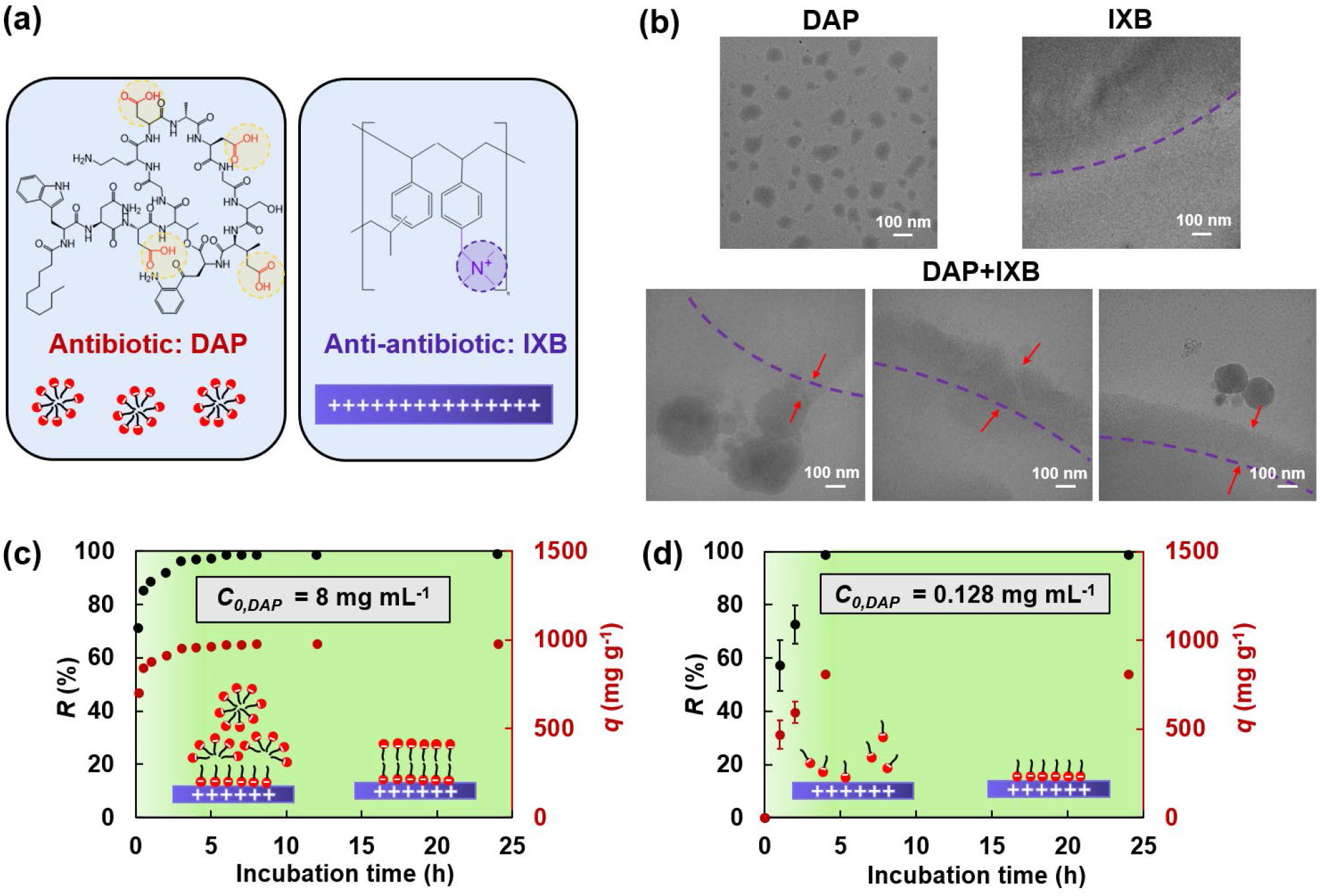
Kinetics of IXB-mediated DAP removal. **(a)** Chemical structure of the antibiotic (DAP) and the anti-antibiotic (IXB) **(b)** Cryo-TEM images of DAP, IXB, and DAP-IXB complex. Kinetics of DAP removal percentage (*R*) and capacity (*q*) of IXB at DAP concentrations of **(c)** 8 mg mL^-1^ or **(d)** 0.128 mg mL^-1^. The CMC of DAP is 0.147 mg mL^-1^. The insets schematically show the adsorption of DAP micelles (panel c, DAP concentration > CMC) or molecules (panel b, DAP concentration < CMC) to the IXB.

The IXB-mediated DAP removal was conducted by incubating IXB with DAP in Milli-Q water. To investigate the time scale of DAP removal at initial DAP concentrations above or below CMC, the effect of incubation time on the removal percentage (*R%*) and capacity (*q*) of IXB was studied, as presented in Figure 1. Figure 1c presents *R* and *q* at DAP concentrations above the CMC. Both *R* and *q* significantly increased by increasing the incubation time and reached a plateau after about 4 h. The *q* reached a plateau value of ∼ 1000 mg g^-1^ after 4 h, i.e., the maximum removal capacity (*q_e_*) equivalent to 100% DAP removal. Similarly, at DAP concentrations below the CMC (Figure 1d), *R* and *q* increased by increasing the incubation time within 4 h and reached their plateau values of 100% and ∼ 800 mg g^-1^, respectively, after 4 h. No significant difference between the IXB saturation time below and above CMC was observed. Therefore, the required time (∼ 4 h) to reach the maximum *q_e_* may not be attributed to the DAP fusion and SLB formation. We hypothesize that the time-dependent adsorption of DAP is a result of molecular diffusion into the IXB pores. To test this hypothesis, we mathematically model the removal of DAP in the next section.

### 3.2 Mathematical and experimental modeling of IXB-mediated DAP removal

Since IXB is a porous, swollen polymer resin,^25^ we considered the dynamic diffusion-adsorption of DAP molecules to model the process. We assumed that the IXB is spherical with no tortuosity. In addition, the DAP mass transfer resistance from the bulk solution to the outer IXB surface is negligible since the solution is well mixed. Accordingly, the bulk DAP concentration (*C*) is at a time-dependent equilibrium at any radial (*r*) position inside the IXB. Thus, the system can be modeled based on the unsteady-state diffusion-adsorption mass balance, prescribed by Eqn. 4:

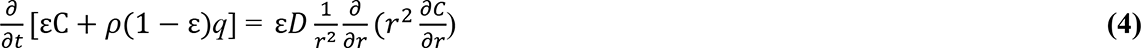

in which the time change of DAP bulk concentration and adsorbed DAP are balanced by the DAP diffusion in the spherical coordinate system. In Eqn. 4, ε denotes the porosity of IXB (0.4),^31^ *ρ* is the IXB density (1.1 kg m^-3^),^25^ *q* denotes the DAP removal capacity at time *t*, and *D* denotes the DAP bulk diffusion coefficient (1.96×10^-10^ m^2^ s^-1^).^26^ To solve Eqn. 4, fractional coverage (θ), defined as the ratio of *q* at time *t* to the equilibrium (maximum) removal capacity (*q_e_*). The initial condition and bounday conditions are listed below.

#### Intial condition

At *t* = 0 and *r* = *R*, *C* = *C*_0_ (at the beginning of adsorption, the DAP concentration on the surface of IXB is equal to the DAP bulk concentration);

#### Boundary condition 1

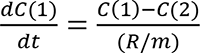 (time change of bulk DAP concentration is equal to the DAP diffusion from the surface to the center);

#### Boundary condition 2

*C*(*m*) = *C*(*m*-1) (symmetry at the adsorbent center).

Here, *R* denotes the IXB particle radius (10.2 ± 2.5 μm, measured by SEM image analysis), and *m* is the number of discretized points in the *r* direction (schematized in Figure 2a). The relationship between the *θ* and the rate contants of adsorption (*k_ads_*) and desorption (*k_des_*) can be expressed as follows:

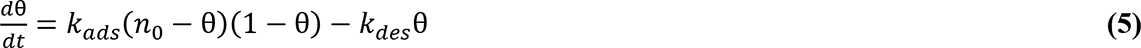

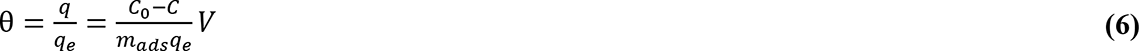

**Figure 2.**
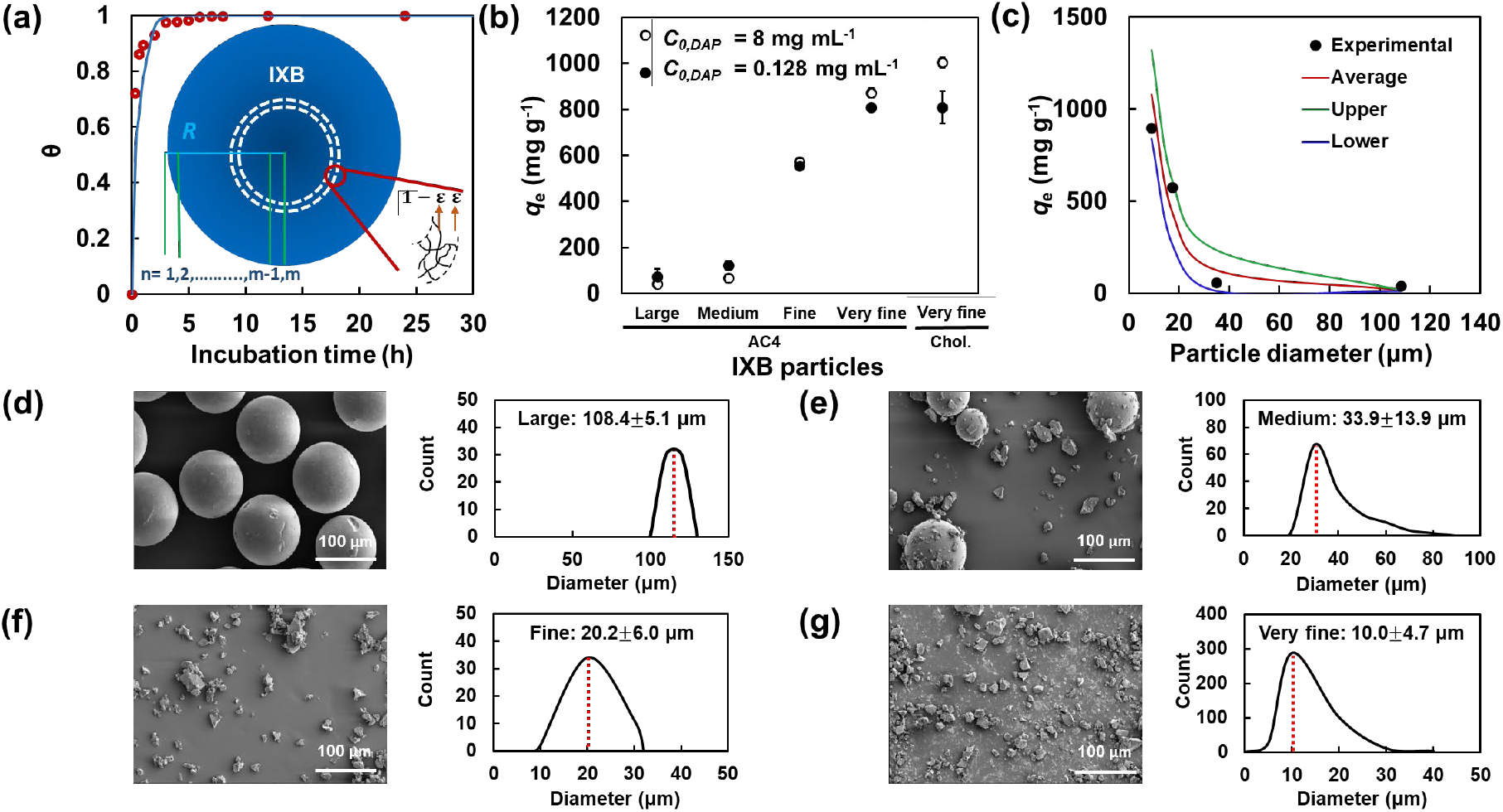
Effect of IXB particle size on DAP removal capacity. **(a)** Fractional coverage of IXB with DAP (8 mg mL^-1^) versus incubation time based on the experimental data and theoretical predictions using Eqns. 4-6 (*R*^2^ = 0.94). **(b)** DAP removal capacity of different sizes of AC4 IXB (incubated for 4 h with a DAP solution, concentration = 8 mg mL^-1^, higher than the CMC, or 0.128 mg mL^-1^, lower than the CMC) compared with the cholestyramine IXB. **(c)** DAP removal capacity of different sizes of AC4 incubated for 4 h with a DAP solution (concentration = 8 mg mL^-1^) based on the experimental data and theoretical predictions via solving Eqns. 4-6 (*R*^2^ = 0.92 for the average, *R*^2^ = 0.90 for the lower limit, and *R*^2^ = 0.87 for the upper limit). SEM images and size distribution of AC4 IXB particles, including **(d)** large particles, **(e)** medium particles, **(f)** fine particles, or **(g)** very fine particles.

Eqns. 4-6 with the initial condition and boundary conditions were solved numerically in Matlab (version R2021a) by converting them into *m* sets of ordinary differential equations (ODEs)^27^ using central finite difference for spatial derivatives (methods of lines, MOL).^28^ Figure 2a shows the time change of IXB fractional coverage calculated by fitting the experimental data with *m* = 30,000 and adjusting *k_ads_*. The plot of surface coverage versus incubation time with the best fit (*R*^2^) resulted in *k_ads_* = 400 s^-1^, corresponding to an adsorption time constant (1/*k_ads_*) of 0.025 s, which proves that the DAP adsorption on IXB is instantaneous. Such a small time constant is in accordance with the DAP SLB formation time scale (<10 s) on negatively charged lipid bilayers, mimicking cell membrane, measured by the quartz crystal microbalance with dissipation,^29^ high-speed atomic force microscopy,^30^ and molecular dynamics.^31^ Therefore, electrostatically driven DAP SLB formation on the IXB may be considered as a nearly instantaneous process. Accordingly, the time (4 h) required to reach the maximum DAP removal capacity is attributed to a diffusion-controlled process. Diffusion of DAP into the IXB may be a result of contact-mediated deformation and de-assembly of otherwise self-assembled DAP molecules. ATR-FTIR spectra of DAP, IXB, and DAP-IXB complex were acquired to confirm the key functional groups involved in the removal process (**Figure S4**). The peaks at 3280 cm^-1^ of DAP ^32^ and 3022 cm^-1^ of IXB spectra were associated with the O-H stretching of carboxylic acid groups and the C-H stretching of quaternary ammonium groups, respectively. The shift of C-H stretching peak to 3034 cm^-1^ in the spectrum DAP-IXB complex, and its broadness may be a result of quaternary ammonium groups-carboxylate groups complex formation. The peak at 3060 cm^-1^ of DAP spectrum was attributed to C-H stretching while the peak at 3357 cm^-1^ of IXB spectrum and the peak at 3335 cm^-1^ of DAP-IXB complex spectrum were both attributed to O-H stretching arising from moisture in the samples.

To further examine the effect of diffusion, other IXBs (AC4) with the same chemical structure as cholestyramine but varying particle diameters were used. The DAP removal capacity at a fixed incubation time of 4 h was measured for large (diameter = 108.4 ± 5.1 μm), medium (diameter = 33.9 ± 13.9 μm), fine (diameter = 20.2 ± 6.0 μm), and very fine (diameter = 10.0 ± 4.7 μm) AC4 particles. Figure 2b shows the DAP removal capacity of AC4 particles with varying diameters after 4 h of incubation compared with the cholestyramine IXB. The removal capacity at initial DAP concentrations higher or lower than CMC decreased by increasing the particle size. To understand the effect of particle size on DAP removal capacity, theoretical *q_e_* based on the solution of Eqns. 4-6 for each particle size (average, upper limit, and lower limit) was calculated and compared with the experimental *q_e_* in Figure 2c. The theoretical *q_e_* decreased by increasing the particle size, matching the experiments, which supports the diffusion-controlled mechanism of DAP removal. The upper limit of the particle diameters resulted in the lower theoretical value of *q_e_*, whereas the lower limit of the particle diameters yielded the upper theoretical value of *q_e_*. The SEM images and size distribution of AC4 particles are shown in Figures 2d-g. The experimental *q_e_* attaining values between the lower and average values of model predictions were associated with the particle diameter distribution skewed from the average to the upper limit of particle diameters (e.g., for the very fine particles, Figure 2g). Together, all of this is consistent with diffusion-controlled DAP removal.

### 3.3 Effect of initial DAP concentration on the equilibrium DAP removal capacity of IXB

The effect of initial DAP concentration on the equilibrium *q_e_* of cholestyramine IXB was studied by incubating the IXB in DAP solutions of 1-20 mg mL^-1^ and measuring DAP concentration in the supernatant after 4 h. Figure 3a shows DAP solutions in Milli-Q water with varying initial concentrations before and after the IXB incubation. The DAP solutions had a uniform yellow color prior to contacting the IXB, which was more distinguishable at higher concentrations, and after contacting the IXB, yellow precipitates were observed at the bottom of the vials. Figure 3b presents the DAP removal percentage and capacity of cholestyramine IXB at varying initial DAP concentrations. The DAP removal percentage was near 100 % when the initial DAP concentrations were below 8 mg mL^-1^. The stoichiometric ratios of quaternary ammonium groups of IXB (mol) to carboxylate groups of DAP (mol) range from 1:0.3125 to 1:2.5 at DAP concentrations ranging from 1 mg mL^-1^ to 8 mg mL^-1^. By increasing the initial DAP concentration beyond 8 mg mL^-1^, DAP removal percentage decreased because the active binding sites of IXB are saturated. The DAP removal capacity of IXB was increased by increasing the initial DAP concentration from 1 mg mL^-1^ to 12 mg mL^-1^ and reached a plateau of ∼ 1250 mg g^-1^ at higher initial DAP concentrations. The maximum DAP removal capacity is ∼200 % higher than the calculated theoretical value based on the charge stoichiometric ratio (i.e., 1 mmol g^-1^ of IXB ammonium groups adsorbs 1 mmol g^-1^ of DAP carboxylate groups, corresponding to 0.25 mmol g^-1^ or 406 mg g^-1^ of DAP). The supra-stoichiometric DAP removal may be a result of DAP self-assembly or SLB formation.

**Figure 3.**
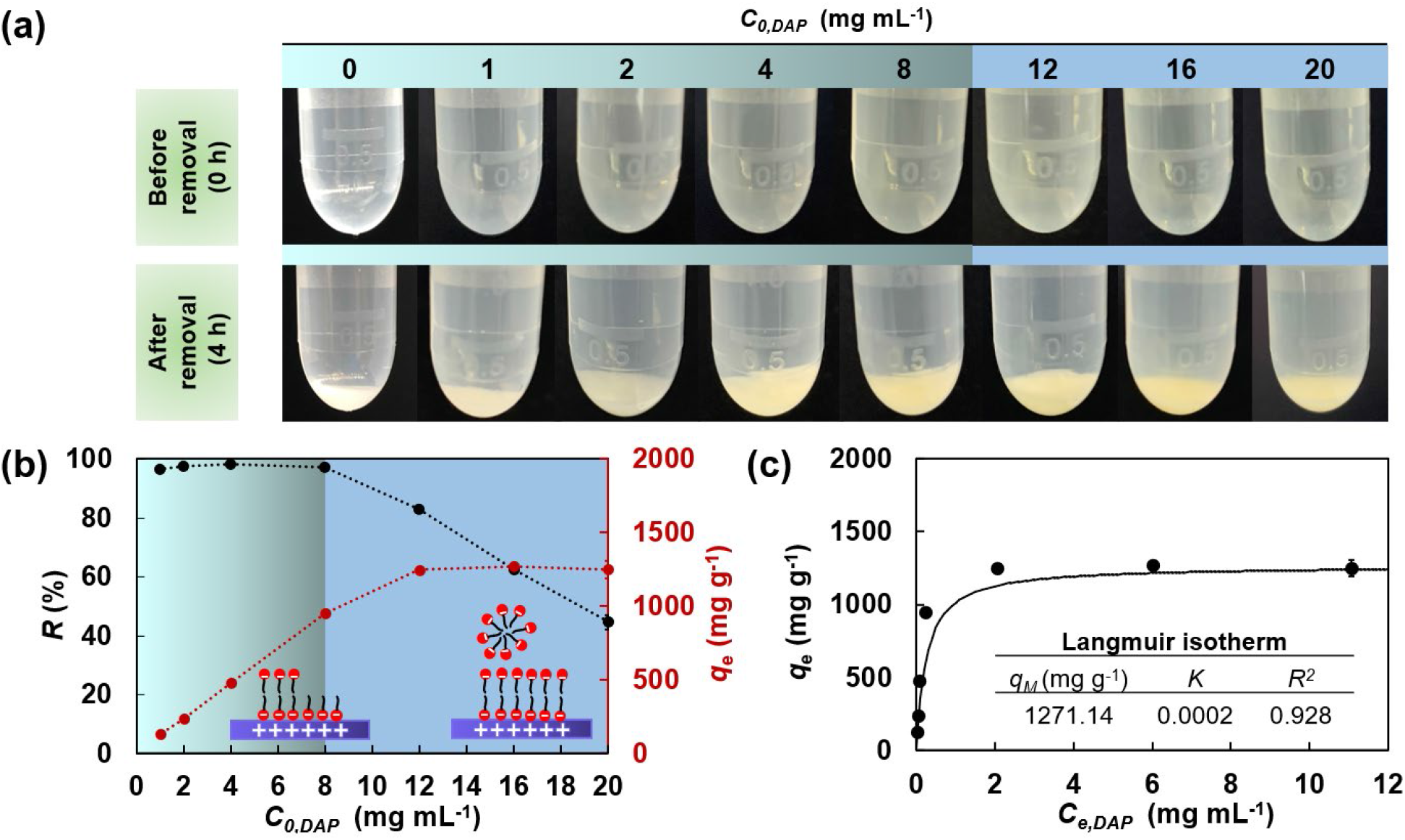
IXB-mediated DAP equilibrium adsorption. **(a)** Photos of DAP solutions with varying DAP concentrations before and after contact with the cholestyramine IXB. **(b)** Effect of initial DAP concentrations on the removal percentage (*R*) and capacity (*q_e_*) of IXB after 4 h of incubation. **(c)** DAP removal capacity of IXB versus equilibrium DAP concentrations fitted with the Langmuir isotherm model. The isotherm parameters are summarized in the inset table.

To understand the adsorption isotherm, Figure 3c presents the equilibrium removal capacity of IXB at varying equilibrium DAP concentrations (i.e., adsorption isotherm). The removal capacity increased as equilibrium DAP concentration increased to 2 mg mL^-1^, reaching a plateau of ∼1270 mg g^-1^. The Langmuir isotherm was fitted to the data using Eqn. 7 ^33^:

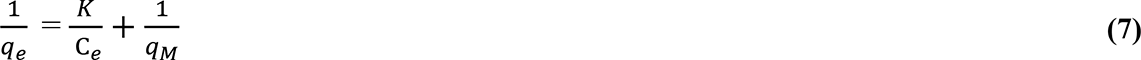

in which *q_e_* is the removal capacity (mg g^-1^) at equilibrium, *q_M._* is the maximum removal capacity (mg g^-1^), and *K* is the adsorption equilibrium constant. Following the Langmuir fitting, the *q_M_* and *K* were ∼1270 mg mL^-1^ and 0.0002, respectively. Although the experimental data were well fitted with the Langmuir isotherm (*R^2^* = 0.928), it should be noted that the adsorption may not simply be a monolayer coverage as a result of DAP self-assembly/SLB formation. The sharp increase of removal capacity at equilibrium DAP concentrations lower than 2 mg mL^-1^ may be associated with the formation of hemimicelles (monolayer adsorption) or admicelles (bilayer adsorption) due to the electrostatic interaction between DAP with the IXB.

### 3.4 Effects of pH and ionic strength on IXB-DAP interactions

To investigate the effect of electrostatic interactions on the cholestyramine IXB-mediated DAP adsorption, pH and ionic strength were systematically altered. Figure 4a shows the chemical structure of DAP, the pK_a_ values of its major functional groups, and its net charge based on ionization states at varying pH.^34^. At pH ranging from 5 to 12 wherein the net charge of DAP is around −4, the DAP removal capacity of IXB remained around 1000 mg g^-1^. Decreasing the pH from 5 to 1.5 decreased the removal capacity. At pH < 5, the carboxyl groups become partially protonated, reducing the number of anionic binding sites of DAP that would otherwise interact with the cationic IXB. These findings imply that the DAP adsorption to IXB is regulated by electrostatic attraction. Figure 4c shows the ζ-potential of DAP at varying pH. At pH = 1.5, the ζ-potential of DAP was positive; however, a DAP removal capacity of 500 mg g^-1^ was still obtained (Figure 4b). This may be explained by discussing three possibilities: (i) although DAP bears a net positive charge, there are still some local negative charges on it, interacting with the positively charged quaternary ammonium groups of IXB; (ii) The equilibrium adsorption reactions (rxn.) for DAP-IXB is explained as follows:

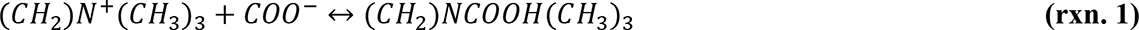

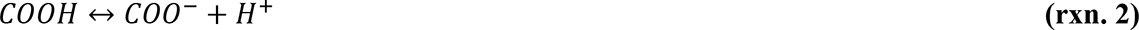

**Figure 4.**
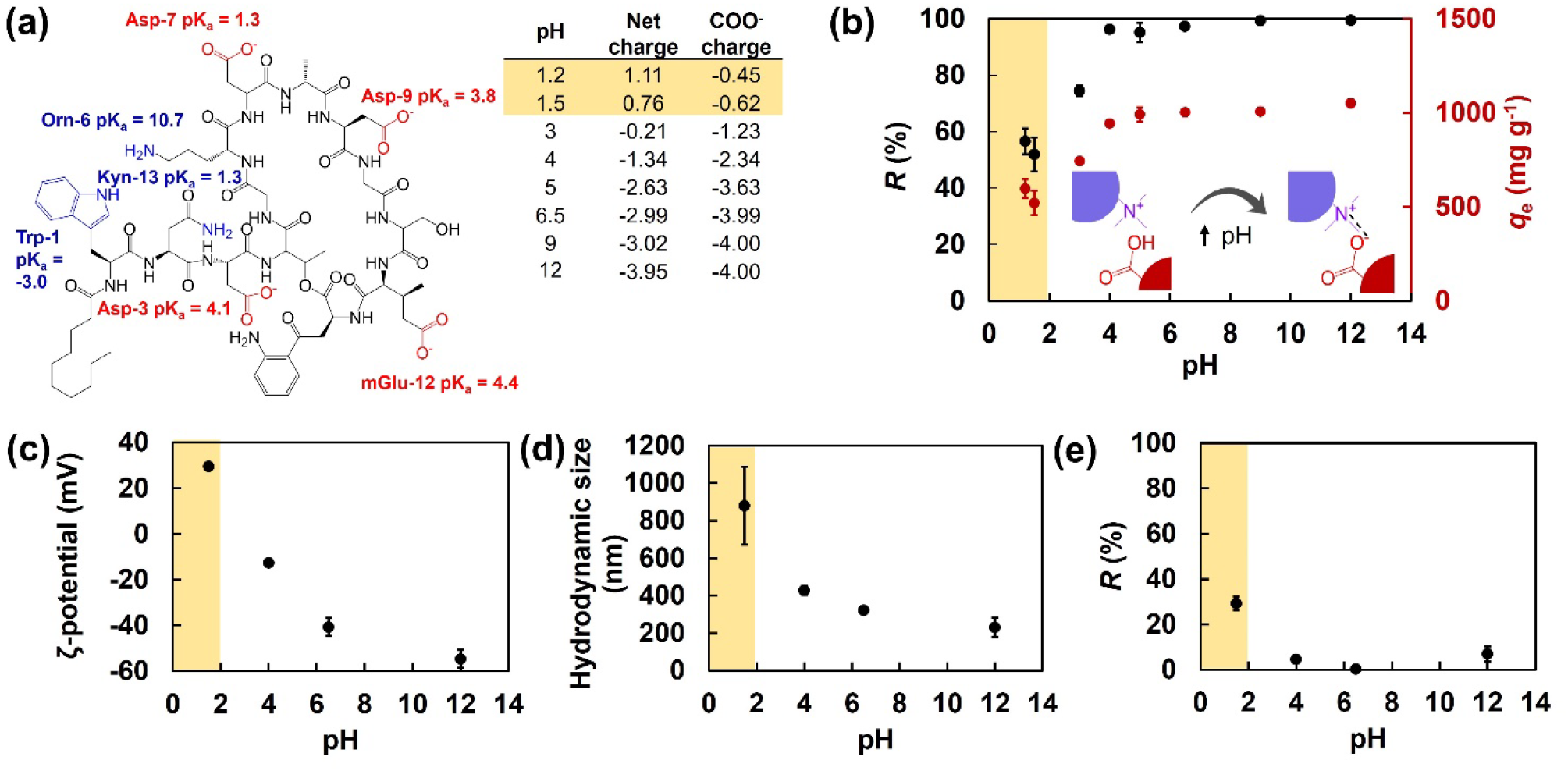
Effect of pH on IXB-mediated DAP adsorption. **(a)** Chemical structure of DAP, the pK_a_ values of its major functional groups, and its carboxylate and net charge at varying pH. **(b)** IXB-mediated DAP (8 mg mL^-1^) removal percentage (*R*) and capacity (*q_e_*) at varying pH. **(c)** ζ-potential and **(d)** hydrodynamic size of DAP at varying pH. **(e)** Removal percentage (*R*) of DAP (8 mg mL^-1^) at varying pH in the absence of IXB. Highlighted pH in yellow indicates a net DAP positive charge and the precipitation of DAP.

Based on the Le Chatelier’s principle,^35^ at low pH (i.e., 1.5), the high proton concentration may shift rxn. 2 to the left; however, the high quatenary ammonium concentration of IXB (8 mM) shifts rxn. 1 to the right, reducing the COO^-^ concentration, which in turn shifts rxn. 2 to the right. Therefore, the concentration of deprotonated carboxylate groups may be higher than the therotical calculation; (iii) DAP may further aggregate and phase separate at highly acidic conditions. To investigate this, DAP hydrodynamic size at varying pH were measured using DLS, as shown in Figure 4d. Decreasing the pH increased the DAP hydrodynamic size from ∼ 200 nm at pH = 6.5 to ∼900 nm at pH=1.5, attesting to DAP aggregation in acidic media. Figure 4e shows the precipitation-mediated removal percetage of DAP (8 mg mL^-1^) at varying pH without IXB. The removal percentage at pH = 1.5 indicates that 30 % of DAP is precipitated out in the absence of IXB. Therefore, the removal capacity at pH = 1.5 is attributed to the DAP self-aggregation and phase separation.

The DAP removal percentage and capacity of IXB in different electrolytes, containing mono-or divalent ions were investigated. Figure 5a shows the *R* and *q_e_* for IXB-mediated DAP removal at varying sodium ion (Na^+^) concentrations. The DAP removal capacity remained almost unchanged, around 1000 mg g^-1^, when the Na^+^ concentration increased from 0 mM to 100 mM. At supraphysiological concentrations of Na^+^, i.e., 200 - 500 mM, the DAP removal percentage and capacity decreased by only ∼ 10% and ∼ 20%, respectively. Figure 5b and Figure 5c show the ζ-potential and hydrodynamic size of DAP at varying Na^+^ concentrations, respectively. Both ζ-potential and hydrodynamic size did not change significantly in the presence of Na^+^. This is likely a result of the DAP charge screening by Na^+^ via decreasing the electrical double layer thickness without comprimsing the surface charge of DAP.^36^

**Figure 5.**
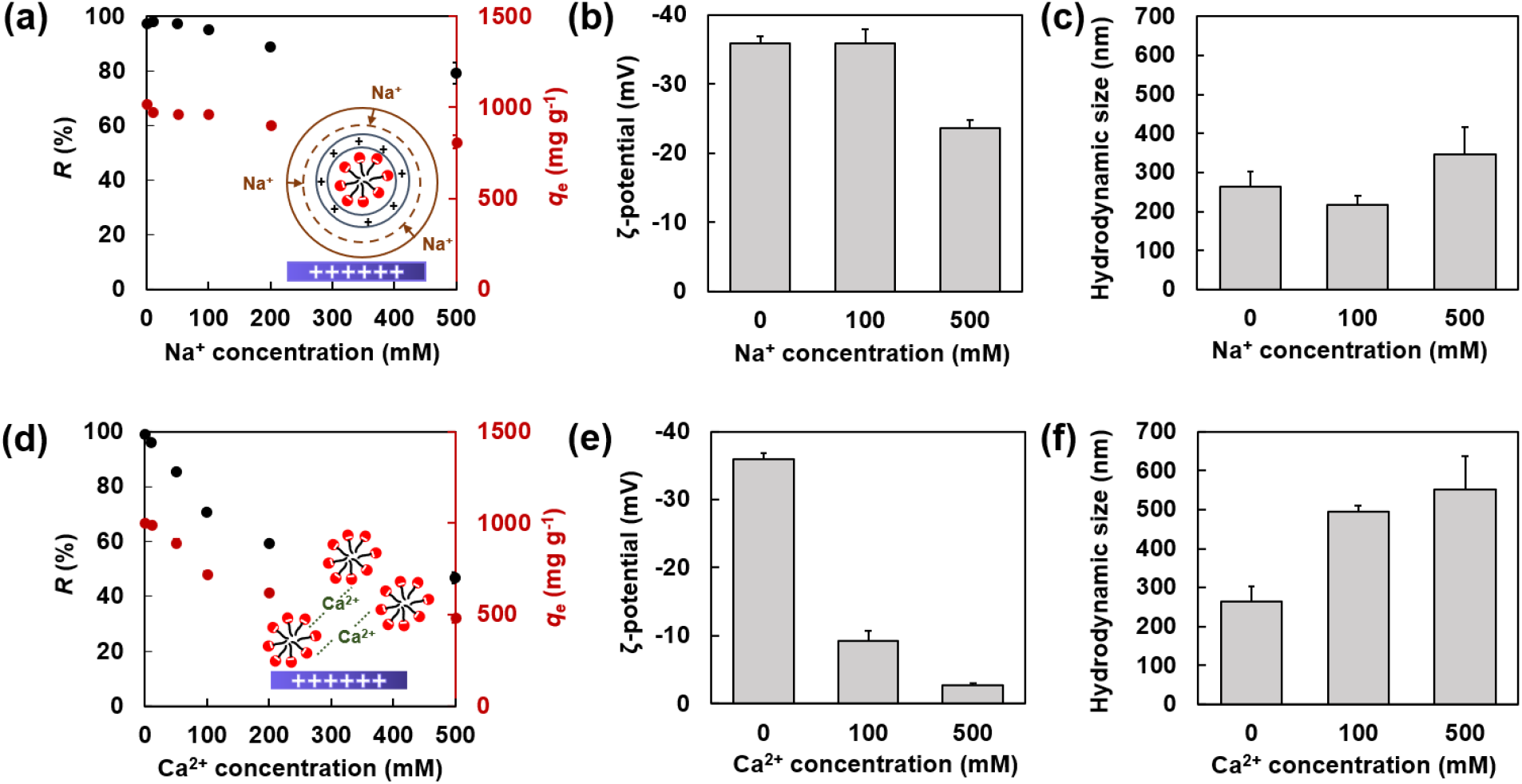
Effects of ionic strength and ion types on IXB-mediated DAP adsorption. **(a)** IXB-mediated DAP (8 mg mL^-1^) removal percentage (*R*) and capacity (*q_e_*) at varying Na^+^ concentrations. The inset shows the Na^+^-mediated shrinkage of electrical double layer of DAP. **(b)** ζ-potential and **(c)** hydrodynamic size of DAP at varying Na^+^ concentrations. **(d)** IXB-mediated DAP (8 mg mL^-1^) removal percentage and capacity at varying Ca^2+^ concentrations. The inset shows the Ca^+^- mediated DAP neutralization and aggregation. **(e)** ζ-potential and **(f)** hydrodynamic size of DAP at varying Ca^2+^ concentrations.

Figure 5d shows the IXB-mediated DAP removal percentage and capacity at varying calcium ion (Ca^2+^) concentrations. The DAP removal capacity decreased more than 50 % when the Ca^2+^ concentration increased from 0 mM to 500 mM, possibly because the divalent ions neutralized the charge (Ca^2+^:COO^-^ = 1 mol : 2 mol).44 Figure 5e and Figure 5f show the ζ-potential and hydrodynamic size of DAP at varying Ca^2+^ concentrations. The ζ-potential of DAP significantly changed from ∼ −36 mV in Milli-Q water to ∼ −10 mV and ∼ −3 mV at 100 and 500 mM of Ca^2+^, respectively. The hydrodynamic size of DAP significantly increased from ∼ 265 nm in Milli-Q water to ∼ 563 nm in the aqueous medium containing 500 mM of Ca^2+^. These results show that Ca^2+^ neutralizes the carboxylate groups of DAP, inducing colloidal aggregation. The effect of Na^+^ and Ca^2+^ concentrations on DAP removal in the absence of IXB was also examined (**Figures S5a** and **S5b**). The DAP removal percentage at 500 mM of Na^+^ concentration without IXB was ∼ 2%, while it was ∼ 10% at 500 mM of Ca^2+^. Accordingly, the majority of DAP removal at high ionic strength is based on the IXB adsorption, and not the DAP precipitation. The pH and ionic strength studies imply that the carboxylate groups of DAP were either protonated or neutralized at low pH or high Ca^2+^ concentrations, decreasing the DAP removal efficacy due to the partial loss of electrostatic interactions. Interestingly, even at such harsh pH or ionic strength conditions, hydrophobic interactions between DAP resulted into phase separation and DAP removal.

### 3.5 Effects of SIF components on IXB-mediated DAP removal

Intestinal fluid contains several types of molecules, such as lipids and bile salts, that may interact with the IXB and affect its DAP removal efficacy. Here, we study the individual effect of SIF components *in vitro* to uncover the competitive DAP removal capability of cholestyramine IXB. Lecithin (PC), one of the zwitterionic phospholipids and also one of the main components of the SIF, is expected to affect the IXB-DAP interactions because the amphiphilic nature of PC may induce additional hydrophobic interactions with DAP without compromising IXB-DAP electrostatic interactions.^38^ PC is also a widespread constituent of the membranes of living cells,^39^ therefore the interactions between DAP and PC would be the biomimetic route of the DAP interactions with cell membrane lipid bilayers.^40, 41^ Figure 6a shows the DAP removal percentage and capacity of IXB at varying PC concentrations ranging from 0.5 mM to 4 mM (physiological concentration range from FaSSIF to FeSSIF). Increasing the concentration of PC slightly increases the DAP removal percentage and capacity of IXB, and at 4 mM of PC, a 10% enhancement in the removal capacity was obtained. Figures 6b and 6c show the ζ-potential and hydrodynamic size of DAP, PC and DAP-PC complex, respectively, at pH ∼ 6.5. Similar to DAP, the ζ-potential of PC was negative (∼ −28 mV) at pH = 6.5, therefore the PC-mediated enhanced DAP removal may not be significantly affected by electrostatic interactions as the anionic PC and DAP compete with each other for the cationic IXB. The hydrodynamic size of PC was ∼ 800 nm while, after the addition of PC to DAP, it reached ∼ 8000 nm, possibly as a result of hydrophobic interactions. Therefore, the enhanced DAP removal capacity of IXB in the presence of PC (∼ 10 %, Figure 6a) may be attributed to the DAP-PC assembly, increasing the number of DAP molecules adsorbed per active site of IXB.

**Figure 6.**
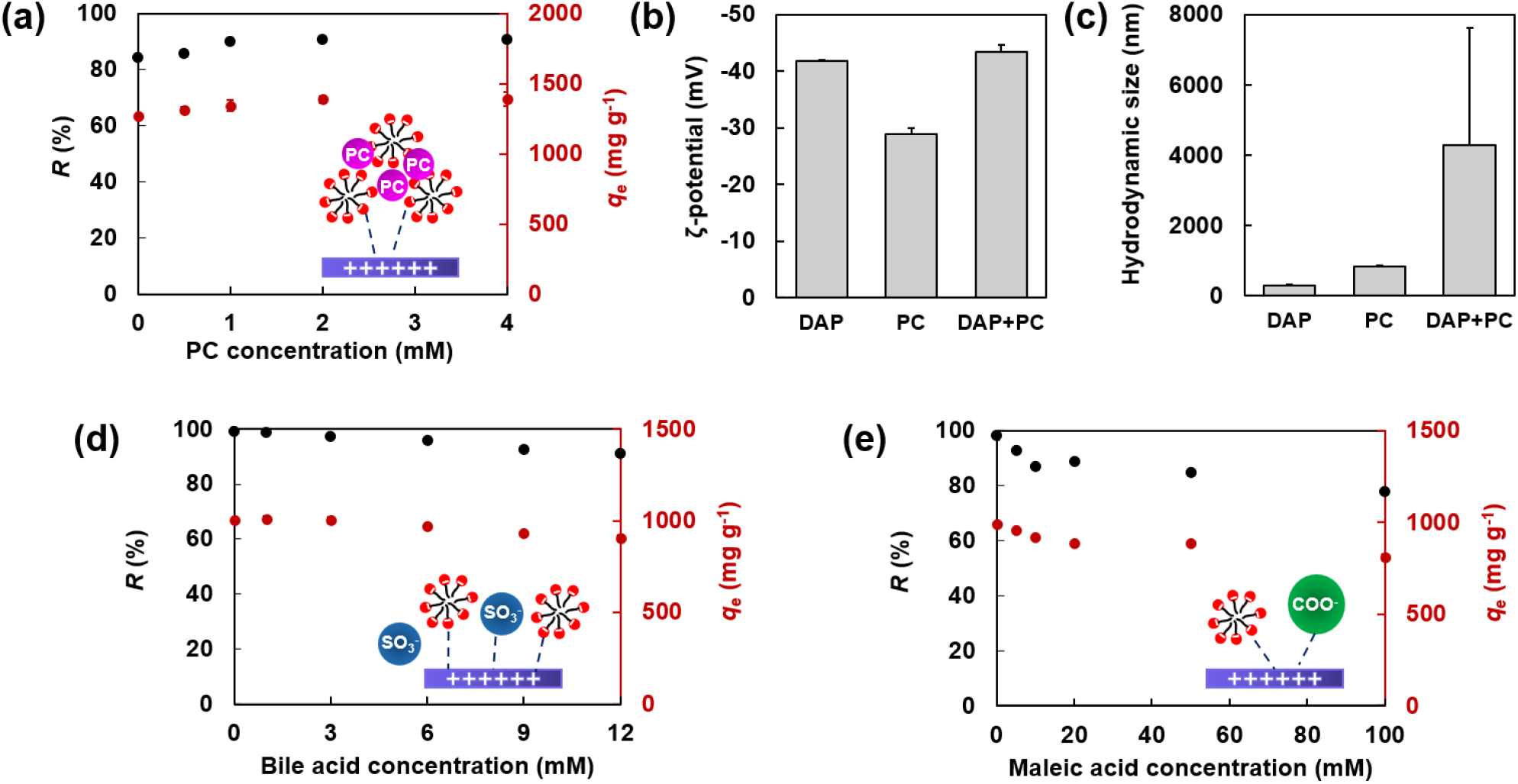
Effects of intestinal fluid components on IXB-mediated DAP adsorption. **(a)** DAP removal percentage (*R*) and capacity (*q_e_*) of IXB at varying PC concentrations (initial DAP concentration = 12 mg mL^-1^). **(b)** ζ-potential and **(c)** hydrodynamic size of DAP, PC, and DAP-PC complex. DAP removal percentage and capacity of IXB at varying **(d)** bile acid concentrations or **(e)** maleic acid concentrations. The insets in panels a, d, and e show the possible interactions among the SIF component, DAP, and IXB. pH was adjusted to 6.5 using NaOH solution in all the measurements.

The DAP removal capacity of IXB was investigated at varying bile acid and maleic acid concentrations. Since IXB has been widely used for sequestering bile acid,^42, 43^ competitive binding of IXB with bile acid and DAP was expected. Figure 6d shows the effect of IXB-mediated DAP removal percentage and capacity at varying bile acid concentrations ranging from 3 mM to 12 mM (physiological concentration range from FaSSIF to FeSSIF). Increasing the concentration of bile acid from 0 to 3 mM does not have any significant effect on the DAP removal, and at higher bile acid concentration, the removal capacity slightly decreases. Despite an equal SO_3_^-^ : COO^-^ molar ratio of the bile acid : DAP (at ∼ 12 mM of bile acid), the DAP removal capacity reduced by less than 10 %. Theoretically, the DAP removal capacity should reduce ∼ 25 % if the electrostatic interactions of bile acid-IXB (SO_3_^-^ - (CH_2_)N^+^(CH_3_)_3_) were the same as the DAP-IXB (COO^-^ - (CH_2_)N^+^(CH_3_)_3_). The bile acid removal capacity of IXB is around 55 mg g^-1^,^44^ which is in consistence with the reduction on DAP removal capacity, as presented in Figure 6d. The effect of bile acid on the ζ-potential and hydrodynamic size of DAP in the absence of IXB are shown in **Figures S6a** and **S6b**, respectively. The ζ-potential and hydrodynamic size of DAP at varying concentrations of bile acid remained almost unchanged, attesting to no significant interactions between them. Self-assembly of bile acid ^45^ also resulted in a hydrodynamic size ∼ 300 nm. Figure 6e shows the effect of IXB-mediated DAP removal capacity at varying maleic acid concentrations ranging from 5 mM to 100 mM (physiological maleic acid concentration in SIF is ∼ 20 mM). When the molar ratio of maleic acid carboxylate groups to the DAP carboxylate groups was identical (at ∼10 mM of maleic acid), the DAP removal capacity of IXB reduced ∼ 7 % compared with the DAP removal capacity of IXB in the absence of maleic acid. This 7 % reduction on the DAP removal capacity was found in Figure 6e in the presence of 10 mM maleic acid.

### 3.6 Effect of fasted state and fed state SIF on IXB-mediated DAP removal

The kinetics of IXB-mediated DAP removal in FaSSIF (**Figure S7a)** and FeSSIF (**Figure S7b)** showed that the removal capacity still reached an equilibrium in 4 h, similar to the removal in Milli-Q water. Figure 7a shows the DAP removal percentage of IXB at varying initial DAP concentrations in the FaSSIF or FeSSIF. The removal percentage was 100 % when the initial DAP concentration was at 1 mg mL^-1^ in both fluids; however, at initial DAP concentrations higher than 2 mg mL^-1^, the percentage of DAP removal in the FeSSIF was always lower than in the FaSSIF. The DAP removal capacity of IXB at varying initial DAP concentrations in the FaSSIF and FeSSIF is presented in Figure 7b. The maximum removal capacities in FaSSIF (∼ 1500 mg g^-1^) and FeSSIF (∼ 1350 mg g^-1^) were higher than in Milli-Q water (∼ 1270 mg g^-1^), which is possibly a result of the additional hydrophobic interactions induced by PC. The lower DAP removal capacity in the FeSSIF compared with the FaSSIF may be explained by the higher bile acid concentration of FeSSIF,^17^ resulting in more competitive adsorption.

**Figure 7.**
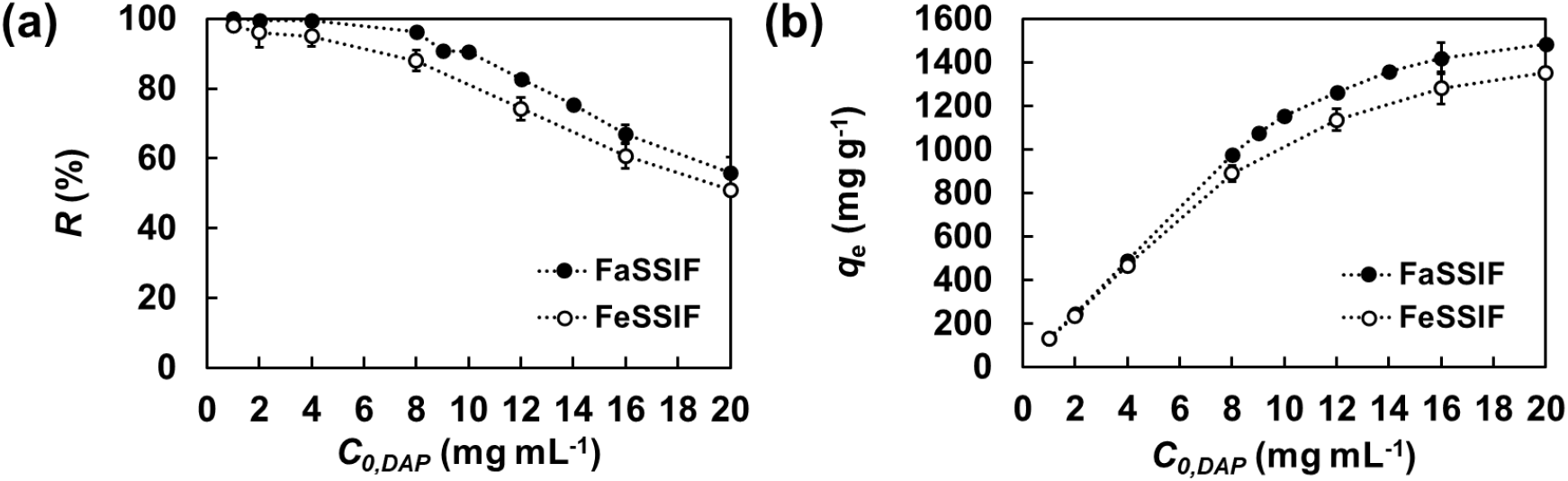
IXB-mediated DAP removal in the SIF. DAP removal (a) percentage (*R*) or (b) capacity (*q_e_*) of IXB at varying initial DAP concentrations in the FaSSIF or FeSSIF.

### 3.7 Antibiotic activity of non-captured DAP

To understand the DAP bioactivity after IXB-mediated adsorption in physiological DAP concentrations, its antimicrobial activity against VRE*fm* at concentrations ranging from 0.25 to 16 μg mL^-1^ was studied in broth microdilution. Figure 8a presents the bacterial densities (OD_600_) following growth in the presence of supernatant collected from DAP with the cholestyramine IXB at different incubation times. The available antibiotic against VRE*fm* reduced with increasing incubation time in the presence of IXB, resulting in an increase in the bacterial density. This is consistent with the time-dependent DAP removal data (Figure 1c,d). Figure 8b presents the bacterial density following growth in the presence of supernatant collected from DAP with varying size of AC4 IXB. The available antibiotic against VRE*fm* was reduced by decreasing the IXB size. The time-dependent and size-dependent DAP removal by IXB in physiological DAP concentrations followed the DAP removal by IXB when initial DAP concentration was higher than CMC, which prove the diffusion-controlled mechanism of DAP removal by IXB.

**Figure 8.**
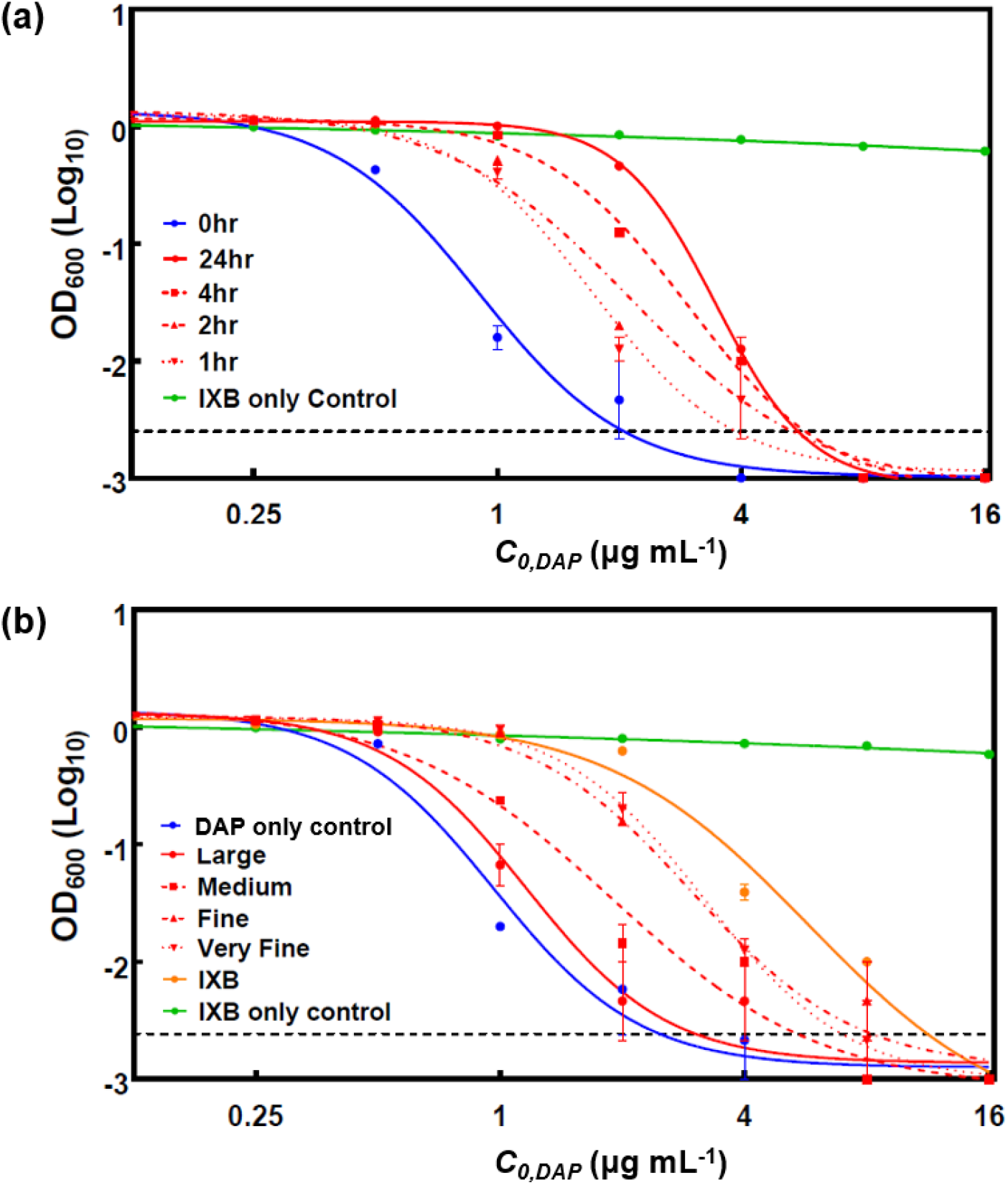
Bacterial densities (OD_600_) for **(a)** DAP with cholestyramine IXB at different incubation time and **(b)** DAP with AC4 IXB with different particle sizes.

## 4. Conclusions

Antibiotics, such as DAP, which enter the gut through biliary excretion, can drive resistance in the opportunistic bacteria in GI tract without therapeutic gain. We previously demonstrated that the oral administration of cholestyramine IXB concurrent with systemic DAP treatment very substantially prevented the up selection and shedding of DAP resistant VRE*fm*.^8, 9^ In this study, we shed light on the mechanism and engineering aspects of IXB-mediated DAP removal. DAP tends to self-assemble into micelles or aggregates in aqueous solutions and re-assemble to form SLBs upon contacting the IXB, which triggers the time-dependent molecular diffusion of DAP inside the IXB pores. The smaller the IXB particle size, the more efficient the DAP removal. As a result of DAP self-assembly, a supra-stoichiometric IXB-mediated DAP removal was obtained. The DAP adsorption by IXB is mainly regulated by electrostatic interactions. DAP undergoes charge neutralization and partial precipitation at a high Ca^2+^ concentration or low pH, which weakens its electrostatic binding to the IXB. Zwitterionic phospholipid molecules (e.g., PC) enhanced the removal capacity of IXB, likely because of hydrophobically-induced DAP-PC aggregate formation, without compromising the electric charges of DAP. Bile acid and maleic acid resulted in some competitive adsorption for DAP but did not significantly disturb the IXB-DAP interactions. The DAP removal efficacy of IXB in the FaSSIF was slightly higher than in the FeSSIF as a result of lower bile acid concentration. This work lays the foundations for optimizing the use of ion exchange sorbents, such as cholestyramine, as adjuvant therapy to prevent daptomycin resistance, as well as designing next generation biomaterials that could combat the emergence of antimicrobial resistance in the GI tract.

## ASSOCIATED CONTENT

### Supporting Information

Calibration lines of DAP UV-vis absorbance at 364 nm; Linear regression analysis of calibration line for DAP in milli-Q water obtained by HPLC; Optical microscope image of cholestyramine IXB; ATR-FTIR spectra of DAP, IXB, and DAP-IXB complex; DAP removal percentage at varying Na^+^ and Ca^2+^ concentrations without using the IXB; ζ-potential and hydrodynamic size of DAP at varying bile acid concentrations; Kinetics of DAP removal by IXB in FaSSIF or FeSSIF.

## AUTHOR INFORMATION

### Corresponding Author

Amir Sheikhi – Department of Chemical Engineering and Department of Biomedical Engineering, The Pennsylvania State University, University Park, PA 16802;

## Acknowledgements

We thank J. Stapleton (Materials Research Institute, Penn State) for connecting us all, A. Zydney for access to his DLS instrument, and the Huck CSL Behring Fermentation Facility for use of the HPLC instrument and A. Sathish for operating it. This work was funded by Penn State (AR, AS) and the Patricia and Stephen Benkovic Research Initiative (AS). Funding from N. Deighton (Huck Institutes of the Life Sciences) made it possible to obtain Figure 1b on the Cryo-EM and Figures 1d and 2b on the HPLC.

