## Supporting Information for "Ion exchange biomaterials to capture daptomycin and prevent resistance evolution in off-target bacterial populations"

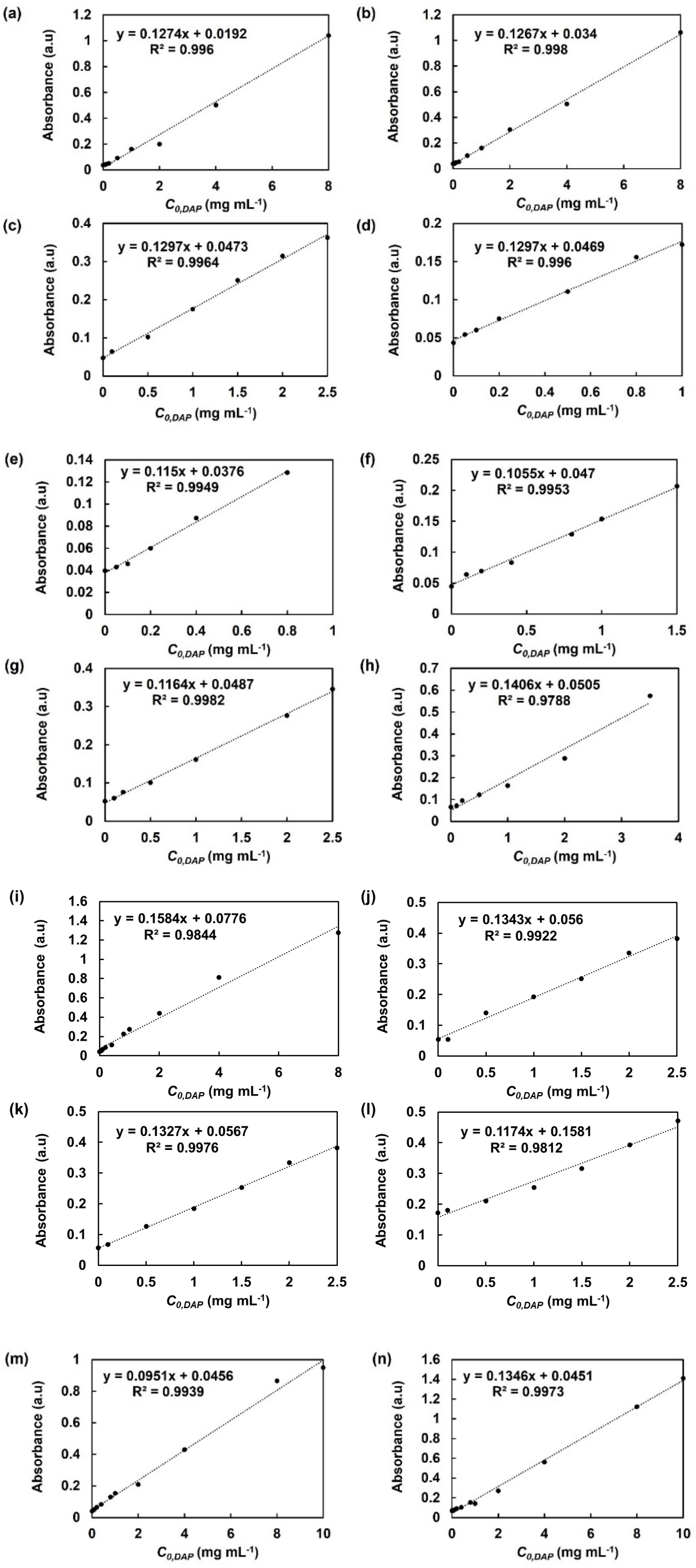


**Figure S1.** Calibration lines of DAP UV-vis absorbance at 364 nm in **(a)** Milli-Q water,

**(b)** NaCl solution (500 mM), **(c)** maleic acid (100 mM), **(d)** bile acid (12 mM), and $CaCl_2$ solutions of **(e)** 10 mM, **(f)** 50 mM, **(g)** 100 mM, **(h)** 200 mM, and **(i)** 500 mM, and lecithin solution of **(j)** 0.5 mM, **(k)** 1 mM, **(l)** 1 mM containing 4 mM of $CaCl_2$, as well as in the **(m)** FaSSIF and **(n)** FeSSIF

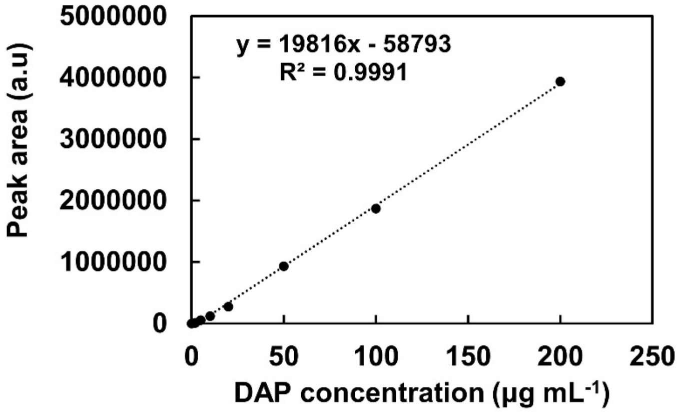


**Figure S2.** Linear regression analysis of calibration line for DAP ranging from 10 µg $mL^{-1}$ to 200 µg $mL^{-1}$ in milli-Q water obtained by HPLC.

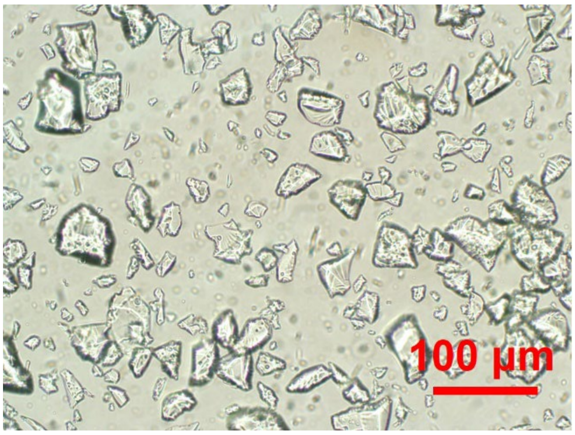


**Figure S3.** Optical microscope image of cholestyramine IXB.

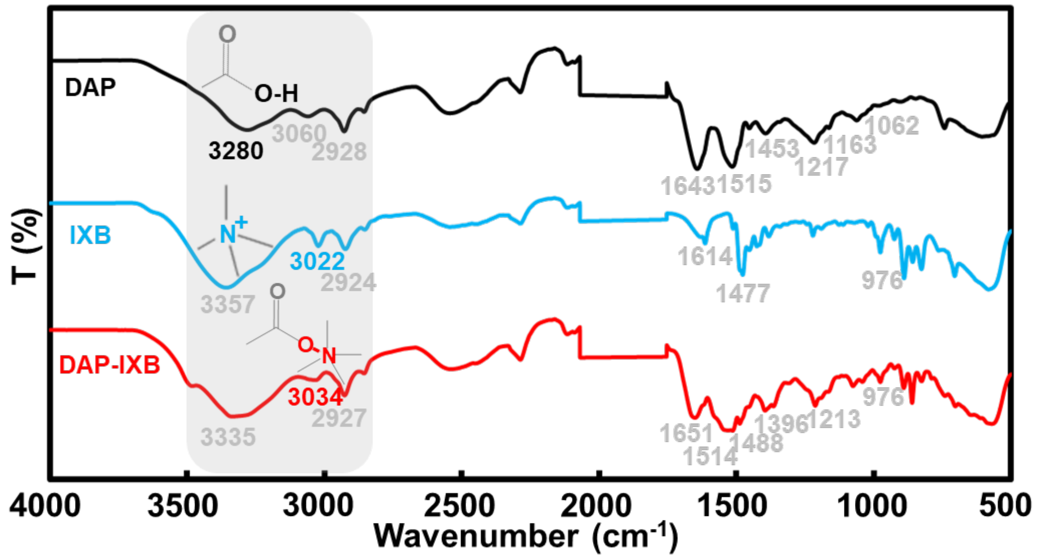


**Figure S4.** ATR-FTIR spectra of DAP, IXB, and DAP-IXB complex.

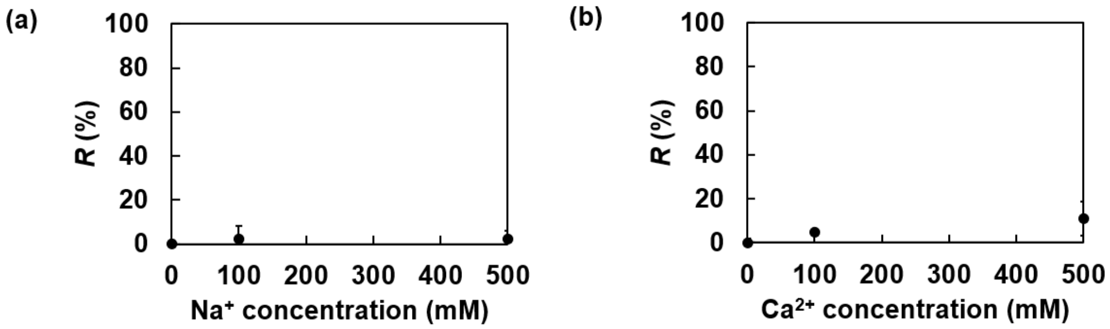


**Figure S5.** DAP (8 mg $mL^{-1}$) removal percentage (precipitation) at varying **(a)** $Na^+$ and **(b)** $Ca^{2+}$ concentrations without using the IXB.

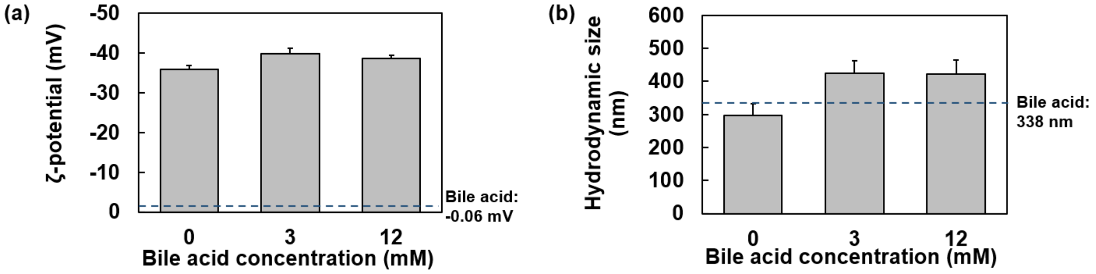


**Figure S6. (a)** ζ-potential and **(b)** hydrodynamic size of DAP at varying bile acid concentrations.

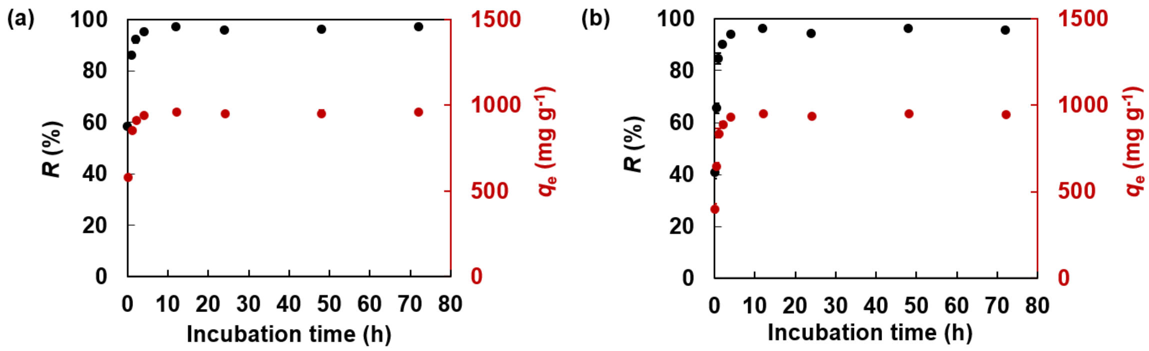


**Figure S7.** Kinetics of DAP (8 mg $mL^{-1}$) removal by IXB (8 mg) in **(a)** FaSSIF or **(b)** FeSSIF.
